# Incorporating host traits into analysis of parasite life history reveals facilitation in coinfection experiments

**DOI:** 10.1101/2025.09.13.675757

**Authors:** Vandana Gupta, Marion Szadkowski, David Carbonell, Benoît Moury, Alison B. Duncan

**Affiliations:** Institut des Sciences de l’Évolution, Université de Montpellier, CNRS, IRD, Montpellier, France; INRAE, Pathologie Végétale, F-84140 Montfavet, France

**Keywords:** virulence, facilitation, multiple infections, *Orthotospovirus tomatomaculae*, *Tetranychus urticae*

## Abstract

Coinfection, when hosts are infected with more than one parasite, can modify life-history traits of both parasites and their host.

We investigated reciprocal interactions between tomato spotted wilt virus (TSWV) and a non-vector spider mite, *Tetranychus urticae*, in coinfections on tomato plants. We compared the number of *T. urticae* daughters and adults and the viral load for two TSWV isolates (France81, which is more virulent, and LYE1137vir) in single and coinfections, and measured host traits (height, fresh weight, chlorophyll content, and number of flowers).

TSWV infection reduced plant height, weight, flowers and chlorophyll content, especially when infected with isolate France81. *T. urticae* only reduced plant height. The more virulent TSWV isolate facilitated spider mites by increasing the number of daughters per unit of plant height, thus potentially enabling equivalent levels of mite transmission as from virus-free plants. In contrast, there was no effect of *T. urticae* on the load of either TSWV isolate.

This work highlights the importance of incorporating measures of host life-history traits when investigating the consequences of coinfection for parasite fitness.

## Introduction

Interactions among parasites in coinfections can modify their life-history traits with consequences for epidemics and evolution [1–4]. Coinfecting parasites may interact via multiple mechanisms, for instance, over shared host resources, via the host immune system and/or effects on host life-history traits, with negative or positive consequences for fitness [2,3,5,6]. The impact of one parasite on another may also change over the duration of an infection: host immunosuppression may facilitate infection with another parasite, which can in turn increase within-host competition [7].

Meta-analyses have found that coinfected hosts more often suffer higher virulence (parasite-induced harm to the host) than corresponding single infections [8,9]. This may be bad for parasites, especially if hosts die before transmission can occur [2,10]. Nevertheless, depending upon transmission mode, the impact of virulence on parasite fitness in coinfected hosts may be more nuanced. For instance, reduced adult emergence in coinfected mosquitoes may increase transmission of the microsporidian *Vavraia culicis* among larvae [11]. Uncovering when and how coinfection impacts both parasite and host life-history traits may enhance understanding about factors promoting transmission and epidemics.

Associations between parasites in coinfection are often non-random [12,13]. Plant viruses that are vectored by arthropods can alter their host’s physiology in ways that attract, and improve host quality for, their vectors [14,15]. These interactions may be mediated via antagonistic cross-talk between different immune pathways [16,17], whereby viruses activate the salicylic acid (SA) immune pathway in plants, which in turn downregulates the jasmonic acid (JA) pathway, preventing an effective immune response against arthropod parasites [18,19]. Viruses can also increase levels of free amino acids circulating in their host plants, which may be a beneficial resource for other co-infecting parasites [20,21]. However, plant viruses exist in complex communities often sharing their host with other non-vector arthropod ectoparasites that can also benefit from changes to plant physiology [20–23] but see [18,24]. In turn, these non-vector arthropods may impact the virus, either by effects on viral load mediated via free-amino acids or the host immune system [21,25], but also as a benefit to the vector as a source of prey [26].

We studied reciprocal interactions between tomato spotted wilt virus (TSWV; *Orthotospovirus tomatomaculae,* family *Tospoviridae*) and *T. urticae* during coinfection on tomato plants (*Solanum lycopersicum*, family *Solanaceae*). Both are generalists with over 1,000 species of host plants [27,28] causing widespread agricultural damage [29,30]. TSWV is vectored by at least nine species of thrips [28], with at least one species, *Frankliniella occidentalis*, producing more offspring when feeding on TSWV-infected plants [31]. TSWV has also been shown to facilitate *T. urticae*, increasing fecundity and the number of offspring becoming adult [20,22,23]. We wanted to explore whether the degree of facilitation for *T. urticae* changed with TSWV isolate and it was linked to viral load (i.e. does more virus equate to more facilitation). It is also not clear whether *T. urticae* can impact TSWV within-host growth. *T. urticae* can induce the JA pathway [32] possibly suppressing the SA pathway via negative immune cross talk [17]. One study found no effect of *T. urticae* on TSWV load when measured at a single time point [20], but we wanted to investigate this further. Finally, we wanted to explore whether interactions between *T. urticae* and TSWV could be mediated via effects on host plant traits. TSWV can infect the whole plant including roots, leaves, petals and stems [33] and probably fruit, with symptoms including chlorotic spots or lesions, systemic necrosis, stunted growth and can ultimately lead to plant death [30,33,34]. Since TSWV inoculation occurred prior to infestation with *T. urticae*, we hypothesised that any increased facilitation due to higher viral load may, in turn, be limited by negative effects on the host. For instance, if higher virulence causes elevated stunting, there may be less plant resources for mites to feed upon.

## Material and Methods

### Biological System

#### Tomato spotted wilt virus

TSWV is a tripartite plant RNA virus with two ambisense (i.e. with both positive and negative polarities) RNAs and one negative-sense RNA strand [30,35]. In our laboratory, we mechanically inoculated plants using leaf extracts from TSWV-infected plants. Briefly, to do this, 1 g of infected leaves was crushed with 4 mL of phosphate buffer (Na_2_HPO_4_.12H_2_O 0.03 M plus 0.2% w/v sodium diethyldithiocarbamate -DIECA) and 90 mg of active charcoal with a pestle and mortar, before adding 75 mg of silicon carbide (Carborundum), which is an abrasive powder. The inoculum was rubbed gently with a finger onto the surface of the first fully expanded tomato plant leaf before being rinsed 15-20 minutes later. Virus-free plants in the experiment were exposed to a simulated inoculation in which virus-free leaves were crushed with the same buffer, mixed with active charcoal and Carborundum and applied to the first fully expanded leaf in the same way as for virus inoculations. This was done to control for any effect of plant wounding during inoculation, which may impact spider mite or plant traits independently of TSWV infection.

For experiments, two virus isolates from the collection at the Plant Pathology Unit, INRAE, Avignon were used (France81 and LYE1137vir). Isolate France81 was collected from a pepper plant (*Capsicum annuum*) in Bouches-du-Rhône (France) in 2008 [36]. LYE1137vir was initially collected from a tomato plant (*S. lycopersicum*) in Drôme (France) in 1996 and subsequently inoculated once onto a pepper (*Capsicum chinense*) accession carrying the *Tsw* resistance gene. No additional passages were performed for either viral isolate, with the isolates being stored as leaf pieces in liquid nitrogen before being transferred to the University of Montpellier in October 2022, following re-inoculation onto tomato plants (cultivar Monalbo) from frozen. At the University of Montpellier, they were maintained by serial transfer every ∼14 days onto 3-week-old tomato plants (cultivar Moneymaker) in the lab for 5 successive infection cycles before use in this experiment.

#### The spider mites

*T. urticae* are haplo-diploid and complete their life cycle in 13-14 days under our laboratory conditions. The *T. urticae* used in this experiment were from an outbred population created by crossing males and females from three different populations collected in Portugal (see [37] for more details). Samples of this population were transferred to University of Montpellier in December 2021 with additional mites transferred and added to the population in November 2022. They were then maintained on three tomato plants (cultivar Moneymaker) in big plastic boxes (dimensions-520 mm x 300 mm x 250 mm) with one plant being changed weekly. Populations were maintained at 25 ± 2°C on a 16: 8 light: dark cycle. To obtain equally aged females for the experiment, 13 groups of 50 mated females were placed together on a cut tomato leaf placed in a jar of water 14 days prior to the experiment to lay eggs. The experiment used the Moneymaker F_1_ hybrid tomato cultivar grown in an isolated arthropod-free environment at 25 ± 3°C.

### The main experiment

#### Interactions between T. urticae and TSWV in coinfection on tomato plants

This experiment measured how coinfection affected tomato plant, *T. urticae* and TSWV life-history traits. There were a total of two spider mite treatments (presence/absence) combined with three virus treatments (isolate France81/isolate LYE1137vir/no virus) (Table 1). Three weeks after sowing, 72 plants were inoculated with the France81 isolate, 72 plants were inoculated with the LYE1137vir isolate and 24 control plants were mock-inoculated (i.e. simulating infection) with a sap extract from healthy tomato leaves as described above. Fourteen days post inoculation (dpi), we transferred 20 adult *T. urticae* females onto a total of 56 plants (24 plants infected with the France81 virus isolate, 24 plants infected with the LYE1137vir virus isolate and 8 control, mock-inoculated plants). The *T. urticae* females were placed in two groups of 10 on a single leaflet of two different leaves (above the inoculated leaf). The mites were isolated on their respective leaflets using a muslin leaf cage (dimensions: 70mm x 90 mm x 10 mm) as in [23] and plants were placed in large plastic boxes (500 x 340 x 400 mm). There were 16 control, mock-inoculated plants without mites. TSWV load was measured at multiple time points (6, 13, 20 and 27 dpi) and required the culling of plants — 12 plants sampled at each time point for each single/coinfection treatment (Table 1). The plants from different treatments were randomised across boxes following the transfer of mites. We counted the numbers of male and female adult *T. urticae* using a binocular microscope on the 28th day following viral inoculation. Note that for plants that were coinfected with TSWV, some leaves were removed on day 27 to measure viral load. Eggs and other developmental stages were not measured as it was not possible to remove and replace the muslin bags without disrupting the mites.

**Table 1:**
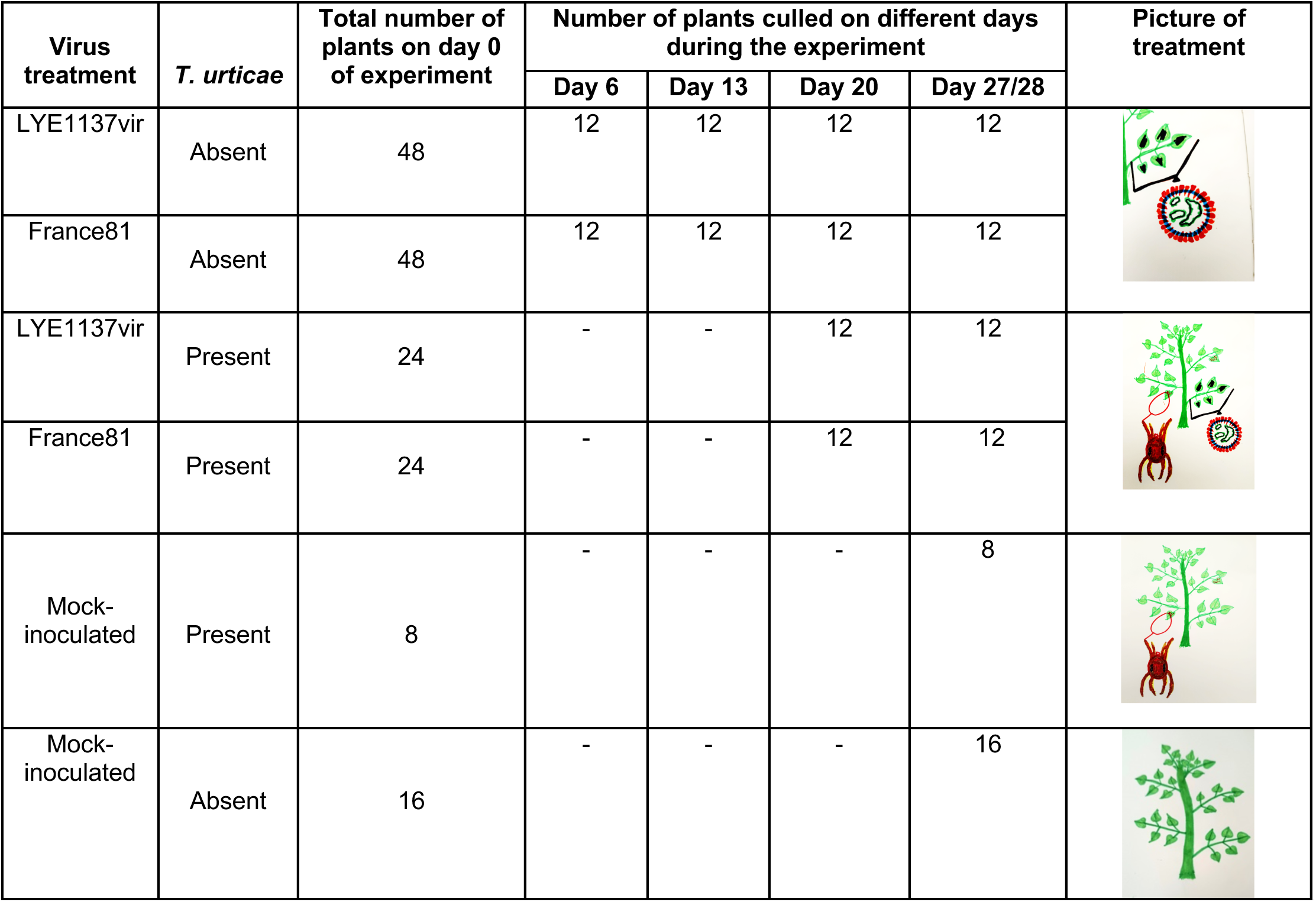
Schematic representation of the experiment design. There were six total categories of treatments comprising single and coinfections with either virus isolate France81 or LYE1137vir, or a mock virus inoculation (3 levels of ‘virus treatment’) in the presence or absence of *T. urticae* (2 levels). Virus load was measured at 4 time points, on days 6, 13, 20 and 27, and required the removal of 12 plants for each virus by mite treatment combination. The column ‘total number of plants on day 0 of the experiment’ denotesthe number of plants allocated to each treatment at the beginning of the experiment. The number of plants removed from the experiment to measure virus load at different time points are marked in the columns labelled Day 6 - 27/28. Day 27/28 is marked as such since leaves were removed from plants to quantify viral load on day 27 and other plant traits were measured on day 28. The last column is a drawing of a tomato plant with TSWV being applied to a leaf (shown in black across the bottom right leaflets) and/or a spider mite to denote groups of 10 (red dots) onto two leaflets above the TSWV or mock - inoculated leaf.

| Virus treatment | *T. urticae* | Total number of plants on day 0 of experiment | Number of plants culled on different days during the experiment |  |  |  | Picture of treatment |
| --- | --- | --- | --- | --- | --- | --- | --- |
|  |  |  | Day 6 | Day 13 | Day 20 | Day 27/28 |  |
| LYE1137vir | Absent | 48 | 12 | 12 | 12 | 12 |  |
| France81 | Absent | 48 | 12 | 12 | 12 | 12 |  |
| LYE1137vir | Present | 24 | - | - | 12 | 12 |  |
| France81 | Present | 24 | - | - | 12 | 12 |  |
| Mock-inoculated | Present | 8 | - | - | - | 8 |  |
| Mock-inoculated | Absent | 16 | - | - | - | 16 |  |

#### Measuring virus load using Quantitative DAS-ELISA (Double Antibody Sandwich - Enzyme-Linked Immunosorbent Assay)

Quantitative DAS-ELISA (qELISA) was used to compare the within-plant load of TSWV between tomato plants. We sampled virus-infected plants on days 6, 13, 20 and 27 by cutting and transferring 1 g of the uppermost leaves into extraction bags (Bioreba, Reinach, Switzerland) that we placed immediately in a fridge at 4°C. The following day, samples were taken to the Plant Pathology Unit (INRAE, Avignon) for qELISA.

We followed the ELISA protocol in Legnani et al. (1995) [38], using TSWV-specific polyclonal antibodies prepared at the Plant Pathology Unit with TSWV isolate LYE51, a tomato isolate collected in 1991 in Berre (Bouches-du-Rhône, France), as an antigen. For each plant, the 1 g of apical non inoculated leaves was ground in a phosphate buffer (1:4 wt/vol). Fivefold dilutions of each plant extract in buffer were tested using DAS-ELISA (from dilution factor 0.2 to 6.4×10^-5^ w/v) and absorbance values were measured at 405 nm (A_405_). The relative virus load of each sample was estimated using curves representing the relationship between the dilution factor and A_405_, within the dilution range where the curves decreased linearly and were parallel among samples, with the help of a home-made program using R version 4.3.2. A common plant sample included on each ELISA plate as a control was used to standardise virus loads across plates, while correcting for differences in enzymatic reaction times.

#### Host plant traits

We measured the height of plants on days 13, 20 and 28 by measuring from just above the cotyledon insertion to the highest branch division. We measured the weight, chlorophyll content, and number of flowers of plants in the different treatments that remained in the experiment 28 dpi. The weight of virus-infected plants that were culled was also measured on days 13 and 20. Chlorophyll content was measured using a single-photon avalanche diode (SPAD) (Konica Minolta, France). This was done by taking three separate measures on the leaf immediately above the leaf that was inoculated with TSWV (lower leaf), and the leaf above the higher leaf with *T. urticae* (upper leaf).

### Statistical analyses

All analyses were done in JMP Student Edition 19.0.4. All models were simplified in a stepwise fashion removing non-significant terms (with p-values > 0.05). Virus isolate and coinfection with *T. urticae* were included in models as fixed factors and day of sampling as a continuous variable. Complete statistical models are shown in tables in the Supplementary materials (Tables S1 – S7). As we were interested in the impact of coinfection on traits, we only include analyses that included both *T. urticae* and TSWV in the main text. See Supplementary Materials for analyses of plant height (Table S1) and viral load (Table S3) across all time points on plants without spider mites. Note that TSWV was not detected in 3/45 plants 21 dpi (one plant inoculated with LYE1137vir with mites and two plants inoculated with LYE1137vir without mites) and 9/45 plants 27 dpi (four plants inoculated with France81 with mites and five plants inoculated with France81 without mites). As all plants from earlier time points were positive for infection, we think viral loads in these plants had declined to levels too low for detection. These plants were not included in analyses that included viral load. We did, however, repeat the analysis for viral load in the presence of mites including these data points, assuming they had a viral load ∼2 orders of magnitude lower than the lowest viral load measured. This did not change the result showing no effect of mites on TSWV load (Table S3). A fungus, we think *Oidium neolycopercisi*, was reported growing on a number of plants with mites 28 dpi. This did not change results, but was included in statistical models for *T. urticae* traits as a random effect. Plants without mites were not checked for fungus.

#### Quantification of host traits

We used a general linear mixed model (GLMM) with a Gaussian error distribution to test the effect of virus treatment, coinfection with *T. urticae*, day of sampling, all treated as fixed factors, and the interactions among them on plant height 20 and 27 dpi. As measures of height were taken from the same mock-inoculated plants at different time points, replicate plant nested within parasite treatments was included in these models as a random term. We used general linear models (GLM) with a Gaussian error structure to measure the effect of virus infection, coinfection with spider mites and their interaction on plant weight and chlorophyll content in both upper and lower leaves 28 dpi. We used a GLM with a Poisson error distribution, corrected for overdispersion, to measure the effect of virus and coinfection with spider mites on the number of flowers on day 28.

#### Virus load using qELISA

We used a GLM with a Gaussian error structure to investigate how TSWV isolate, coinfection with *T. urticae*, day of sampling, all treated as fixed factors, and their interactions impacted virus load.

#### Effect of TSWV on T. urticae life-history traits

We used a GLMM with a negative binomial error structure and log link function to test whether virus isolate and leaf (lower versus higher) and their interaction affected the total number of adult daughters. We tested whether offspring sex ratio (the proportion of male offspring across both leaves) was affected by viral isolate using a GLMM with a binomial error structure and logit link function. Plant identity was included in both models as a random effect, the first to control for multiple measures from the same plant and the second to correct for overdispersion.

As viral infection significantly reduced plant height, we analysed how virus treatment affected the number of offspring per cm of plant height using a GLM with a Gaussian error structure. We used females per cm, rather than including plant height in models, because there was very little overlap in the height of plants from the different treatments. We present results on adult daughters as they are the dispersing sex. Note that analyses for the total number of adult offspring and total offspring per cm of plant height produce similar results (not shown). Three leaflets with 3 or fewer offspring on the leaflet were excluded from analyses (2 because leaflets had died or broken, and one because all juveniles on the leaflet had died).

#### Quantitative relationships between host traits, viral load and number of spider mites

To visualise the covariation among all quantitative host (height, weight, number of flowers and mean chlorophyll content on higher and lower leaves) and parasite traits (number of adult females and log relative viral load), we used principal components analysis (PCA) on standardised data for all virus-inoculated plants. This analysis included plants measured on days 20 and 27, with and without *T. urticae*. Note that measures of flowers, chlorophyll content and *T. urticae* females were only available for day 27. Finally, we used path analysis by extracting standardised beta-coefficients from regression analyses to investigate if there were any relationships between standardised log relative viral load, plant height and the number of adult daughter spider mites on day 27.

## Results

### Effect of virus and spider mite infection on host plant traits

Both virus treatment (F_2, 109_ = 108.84, p < 0.001) and *T. urticae* presence (F_1, 111_ = 12.10, p = 0.0007) significantly reduced plant height (Figure 1). However, there were no significant interactions between virus isolate and mite (virus*mite; F_2, 107_ = 2.80, p = 0.0653, virus*mite*day; F_2, 122_ = 0.32, p = 0.7267) indicating that their effects on height were independent (see Table S1). Overall, plant height increased over the course of the sampling period (F_1, 122_ = 79.56, p < 0.001) (Figure 1). However, an interaction between virus treatment and day of sampling (F_2, 126_ = 5.73, p = 0.0042) showed that the virus isolates affected height differently. Isolate France81 always reduced plant height relative to control plants, but the relative height difference of plants infected with LYE1137vir changed through time, initially being closer to France81-infected plants, but becoming more similar to control plants 28 dpi (Figure 1).

**Figure 1:**
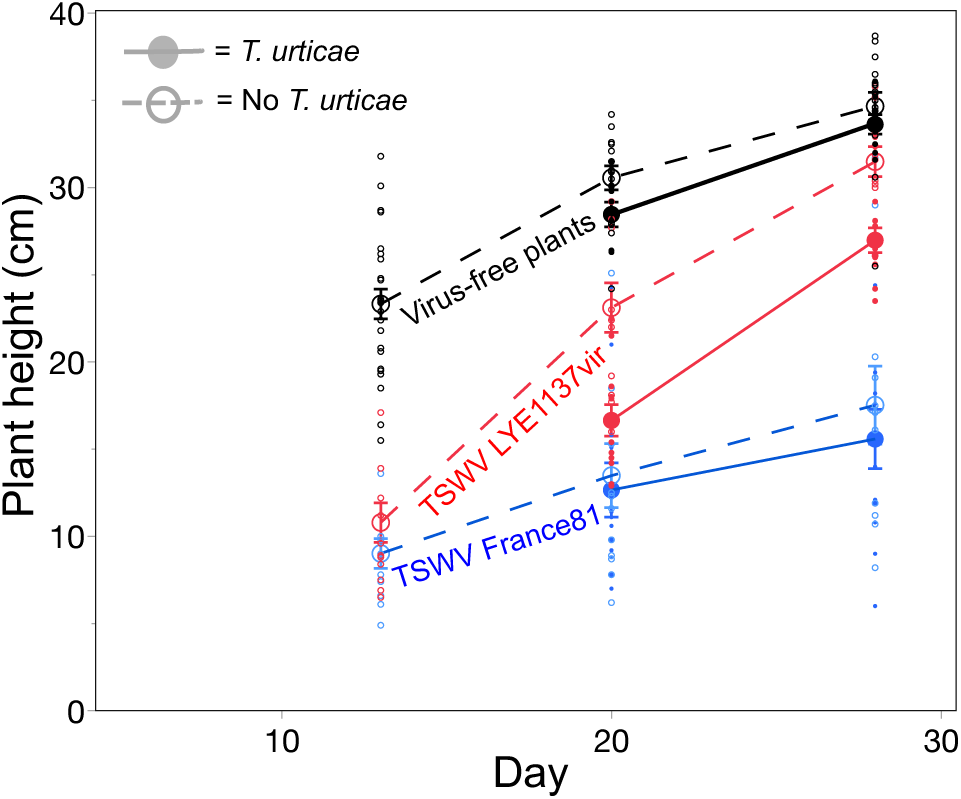
Effect of tomato spotted wilt virus infection through time on mean (± standard error) height of plants inoculated with TSWV isolate France81 (blue), with TSWV isolate LYE1137vir (red), or mock inoculated (black), either infested (closed circles, solid line) or not (open circles, dashed line) with *T. urticae.* Small circles represent individual replicates on the different days.

Virus infection significantly reduced plant weight (**χ**^2^_2_ = 95.86, p < 0.001; Figure S1a), number of flowers (**χ**^2^_2_ = 17.20, p < 0.0001; Figure S1b), and chlorophyll content in upper leaves (**χ**^2^_2_ = 25.31, p < 0.001; Figure S1c). In contrast, *T. urticae* had no effect on these traits (weight; **χ**^2^_1_ = 2.47, p = 0.1160, flowers; **χ**^2^_1_ = 2.10, p = 0.1477, chlorophyll content; **χ**^2^_1_ = 0.004, p = 0.9493, Figure S1). There were no significant interactions between virus infection and *T. urticae* infection for plant weight, number of flowers, or chlorophyll content (Table S2) and no effect of virus (**χ**^2^_2_ = 5.55, p = 0.0623) or *T. urticae* (**χ**^2^_1_ = 1.65, p = 0.1988) on the mean chlorophyll content of lower leaves.

### Virus load using qELISA

There was no effect of coinfection with *T. urticae* on viral load (*T. urticae* main effect: **χ**^2^_1_ = 0.26, p = 0.6105; for all interaction terms with *T. urticae* p > 0.1321, Figure 2; Table S3). However, the France81 isolate had a higher load than LYE1137vir (isolate; **χ**^2^_1_ = 11.82, p < 0.0001) and for both isolates, load declined through time (sampling day; **χ**^2^_1_ = 11.94, p = < <0001).

**Figure 2:**
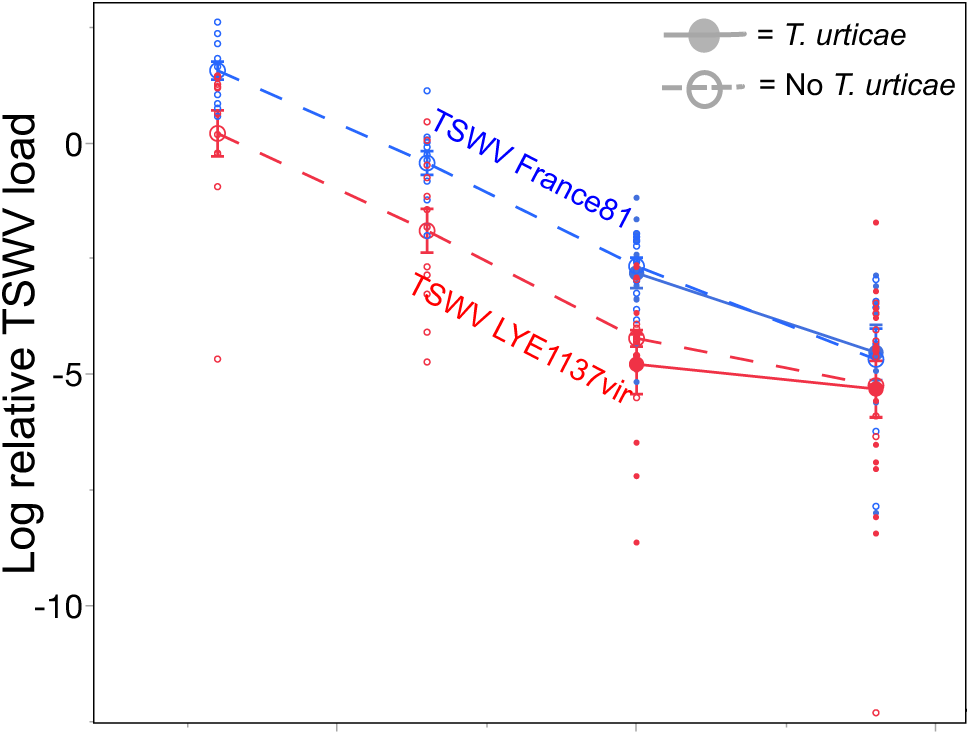
Mean log relative virus load (± standard error) through time for plants inoculated with TSWV isolate France81 (blue circles) or isolate LYE1137vir (red circles), with (closed circles, solid line) or without (open circles, dashed line) the spider mite *T. urticae.* Small circles represent individual replicates.

### Effect of TSWV on T. urticae life-history traits

There were a significantly higher number of daughters per cm of height on plants infected with viral isolate France81 compared to infection with LYE1137vir or control plants (F_2, 29_ = 18.72, p < 0.0001; Figure 3a, Table S4). However, there was no effect of viral isolate on the total number of adult daughters (F_2, 27_ = 0.76, p = 0.4769; Figure 3b) or interaction between leaf and virus treatment (F_2, 27_ = 0.06, p = 0.9396; Figure S2a), despite more daughters on the upper leaf (F_1, 29_ = 4.76, p = 0.0375). Virus treatment had no effect on the offspring sex ratio (F_2, 15_ = 1.08, p = 0.3646; Figure S2c).

**Figure 3:**
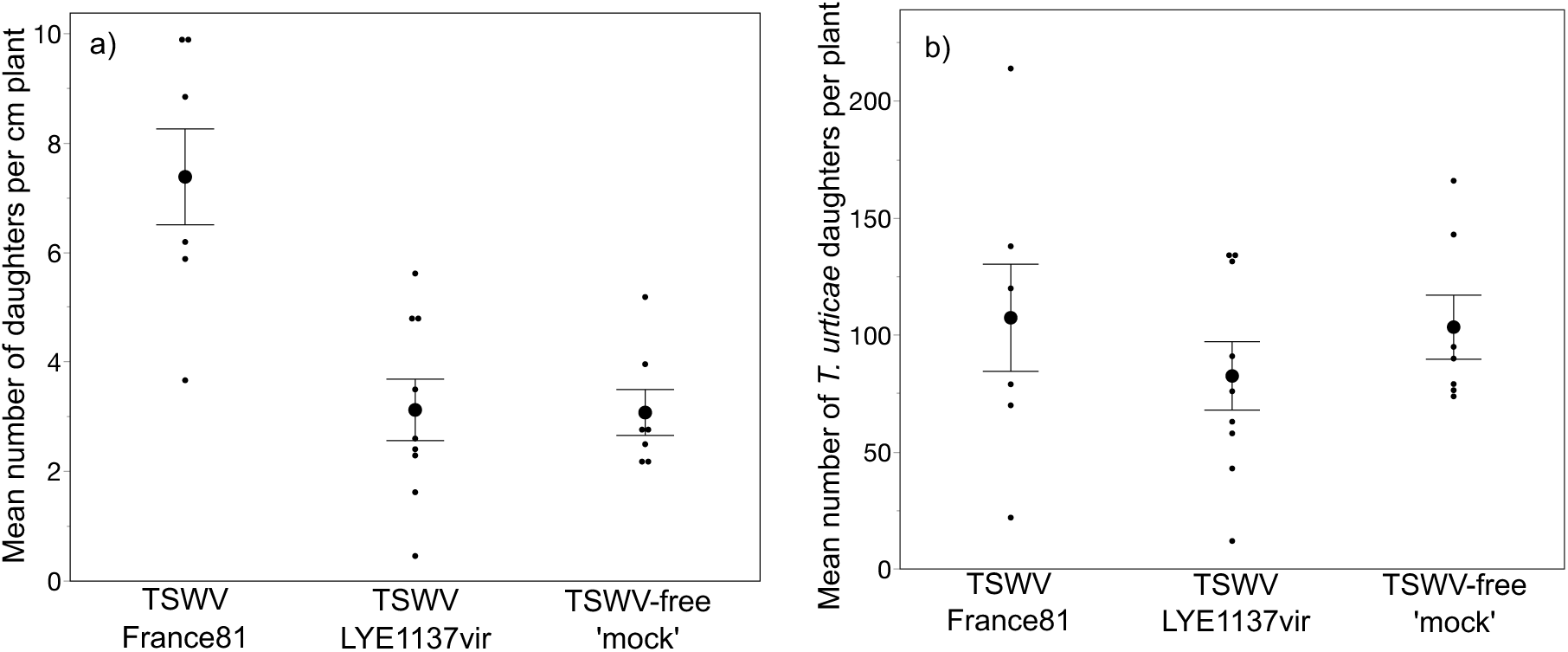
Mean (± standard error) a) total number of *T. urticae* daughters per cm of host height and b) total number of *T. urticae* daughters on plants infected with either TSWV isolate France81 or LYE1137vir or on mock-inoculated plants. Smaller circles represent individual replicates.

### Quantitative relationships between host traits, viral load and number of T. urticae

The PCA showed that axes 1 and 2 contributed 37.4% and 21.7%, respectively, to variation in the data (Figure 4; see Table S5 for the variation explained by the 7 principal components). Axis 1 correlated positively with host plant weight and height and negatively with log relative viral load (Table S6). Axis 2 correlated positively with the number of *T. urticae* females on a plant and lower leaf chlorophyll concentration, but negatively with the upper leaf chlorophyll concentration (Table S6). On day 27, we could not detect a significant quantitative relationship between viral load, plant height and the number of adult daughters using structural equation modelling (standardised beta correlation coefficients and associated p-values for virus load – plant height: -0.038, p = 0.8741; plant height – mite daughters: -0.067, p = 0.8201, virus load – mite daughters: 0.543, p = 0.0841).

**Figure 4:**
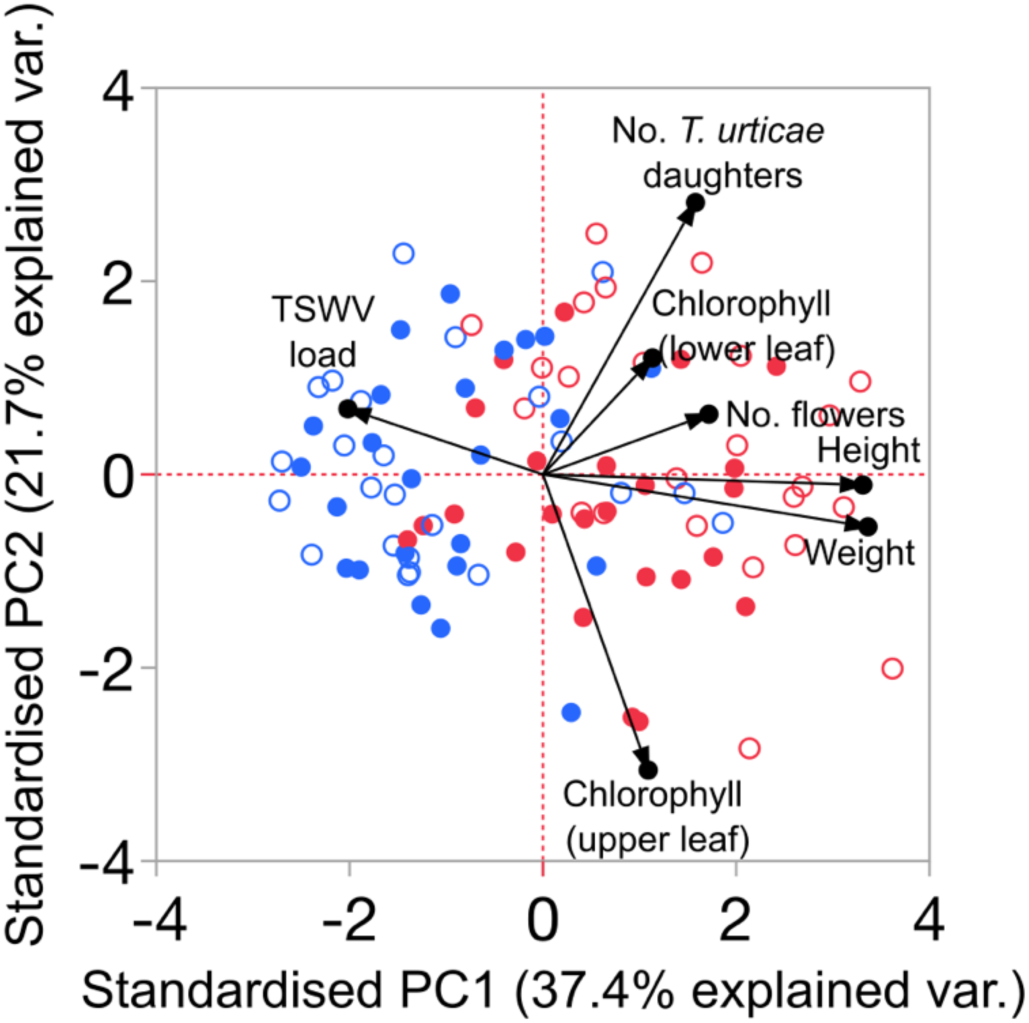
PCA for the five host traits (plant height, weight, number of flowers and mean chlorophyll content in lower and upper leaves), viral load and number of *T. urticae* adult females. Blue circles represent data from tomato plants infected with the France81 TSWV isolate and red circles those infected with the LYE1137vir TSWV isolate. Open and filled symbols represent the absence and presence of *T. urticae*, respectively.

## Discussion

We investigated how coinfection impacted the life-history traits of both *T. urticae* and TSWV, and their tomato host. *T. urticae* had no effect on viral load and, contrary to previous work, TSWV had no effect on the total number of adult *T. urticae* daughters. However, both parasites reduced host height, and it was only when considering this that we could identify an effect of coinfection on *T. urticae*; there were more adult daughters per cm of plant height on tomato plants infected with the most virulent TSWV isolate (France81). Thus, detection of facilitation of *T. urticae* by TSWV required incorporation of host plant traits into measures of parasite life-history.

It seems that, when it has been looked for, coinfections generally induce higher virulence than single infections. One meta-analysis of human parasites reported coinfection to cause greater reductions in host health [9] and another across multiple animal hosts found coinfection to be more virulent, measured as reductions in longevity, than corresponding single infections [8]. This latter study showed that, in coinfection, overall virulence tends to resemble that of the more virulent parasite in the interaction [8]. Our results are consistent with this observation. TSWV was more virulent than *T. urticae*, negatively impacting multiple traits (plant height, chlorophyll content, weight and flowers), especially isolate France81, whereas *T. urticae* only reduced plant height. Further, in coinfection, virulence mostly resembled single TSWV infections, except for plant height for which there was an additive effect of both parasites. Higher virulence in coinfection can reduce parasite fitness via reduced opportunities for transmission if hosts die sooner [2,39,40]. It is plausible, although not observed within the time-frame of our experiment, that the higher virulence in coinfection could in the longer term lead to reduced parasite fitness for both TSWV and *T. urticae*.

In our experiment, *T. urticae* were on plants for a 2-week period, which is too short for all effects of virulence to have established. *T. urticae* have been observed to reduce fruit yield in tomato [41], cucumber [29] and strawberry [42], however, impact in these experiments generally took a few weeks to be observed [29,41,42]. TSWV can also reduce the number and size of tomato fruits [43,44], and consistent with this we found fewer flowers on TSWV-infected plants, especially the more virulent France81 isolate. However, we do not think parasite effects on plant reproduction will impact either parasite in single or coinfection. Although plant viruses can castrate their host [45], which has been associated with increased resources for parasite growth in other systems [46], this is probably not happening here as plants infected with the more virulent isolate, with fewer flowers, were smaller. Reductions in plant height, that were observed in response to both *T. urticae* and TSWV, may be linked to other disease symptoms. In our laboratory, shorter plants generally have a smaller leaf area (unpublished data) and, in another experiment, reduced leaf area caused by cauliflower mosaic virus was positively correlated with host mortality [47].

Coinfection with *T. urticae* did not change virus load on days 20 and 27 for either TSWV isolate, indicating a neutral effect of spider mites on TSWV. This is consistent with previous findings for effects of spider mites on TSWV [20], but is in contrast to other studies which found that non-vector arthropod pests can either increase [25] or decrease [21] plant viral load. The lack of an effect of spider mites on TSWV load could be due to a number of reasons (see [48] for further discussion). Interactions among TSWV and *T. urticae* are likely, in part, mediated via negative immune cross-talk among different branches of the plant immune system, with TSWV inducing the SA pathway [49] and *T. urticae* the JA pathway [32]. One possibility is that there were too few *T. urticae* on the plant to induce the JA pathway sufficiently to suppress the SA pathway. Indeed, induction of genes implicated in the JA pathway can increase in a dose-dependent manner in response to increasing *T. urticae* densities [50]. This may be compounded by asymmetry in hormone feedback, whereby higher levels of JA are required to suppress SA than vice versa [17]. Another possibility is that the spatial distribution of *T. urticae* and TSWV around the plant affected the potential for negative immune cross talk. JA expression can remain local [51] and *T. urticae* were restricted on two leaflets above the site of TSWV inoculation. TSWV, although systemic, takes time to spread within the plant, first being detected in the apical leaves then descending to lower leaves [23]. Any effect of *T. urticae* on TSWV load may require both to be sharing, and measures to be taken from, the same leaf. Elevated levels of circulating free amino acids in virus-infected plants is another mechanism implicated in mediating interactions between viruses and arthropods and is postulated to be the reason non-vector whitefly reduces the load of pepper golden mosaic virus [21]. Low densities and spatial distribution on plants may also have impacted any effect of *T. urticae* on TSWV mediated via free amino acids. Thus, at least at low spider mite densities, sharing the host with this non-vector species does not affect TSWV load.

Contrary to previous work [20,22], including in our laboratory [23], we did not find that TSWV had a positive effect on the total number of *T. urticae* offspring. However, when accounting for plant height, *T. urticae* had ∼4 more adult daughters per cm on plants infected with the more virulent TSWV isolate (France81) than the less virulent isolate (LYE1137vir) or mock-inoculated plants (France81 = 7.38 ± 0.88 SE daughters per cm; LYE1137vir = 3.13 ± 0.56 SE daughters per cm; mock = 3.08 ± 0.42 SE daughters per cm). Previously we had not considered the effect of TSWV or spider mites on virulence or how this may impact parasite fitness. Here, we found that TSWV infection reduced plant height and weight, traits which are positively correlated (Figure 4), and a good proxy for resources available for *T. urticae.* Plants infected with the virulent TSWV France81 isolate were more than 12 cm shorter than uninfected plants on the day of *T. urticae* exposure, and ∼17 cm shorter 27 dpi when *T. urticae* became adult (Figure 1). Thus, it seems that TSWV infection enables *T. urticae* to maintain equivalent offspring numbers when fewer resources are available. In a study with *T. evansi,* another spider mite species, we also only observed a positive effect of TSWV on the number of adult daughters when accounting for plant height when parasite exposure was simultaneous [48].

Despite plants infected with France81 having higher viral loads, we did not find a relationship between TSWV load and the number of *T. urticae* female adult offspring on 28 dpi. This might be because viral load on 28 dpi was too low giving us little power to detect a relationship. In fact, it was not possible to detect virus in 9/48 plants that had been inoculated with the virus 28 dpi, whereas virus was detected in all plants 6 dpi, 13 dpi and 45/48 plants 20 dpi. As nearly all plants prior to 28 dpi were positive, we think that virus levels in these plants had declined to levels below that possible for detection. For both TSWV isolates, the plant systemic load was highest 6 dpi before declining through time until 27 dpi. Declines through time were observed for cucumber mosaic virus, pepper mild mottle virus and pepper mottle virus in pepper plants [52], as well as the daily rate at which new cells became infected for tobacco etch virus in tobacco [53], with a hump-shaped curve describing cells simultaneously infected with different viral variants of cauliflower mosaic virus [54]. An increase followed by a decline in relative viral load may arise due to the plant immune system kicking in and limiting TSWV growth [53,54]. Lagged differences in viral load and spider mite densities might have revealed quantitative relationships between traits, but could not be explored here due to culling of plants for measures of viral load.

Although our measures of parasite fitness are proxies, we do think they provide good estimates for actual transmission. The quantitative DAS-ELISA used to estimate systemic viral load is based on polyclonal antibodies that bind multiple virion epitopes, mostly of the nucleocapsid protein and the glycoproteins of the virus envelope and is therefore representative of the total virion load in the sampled leaf tissues. Further, a positive correlation was found between TSWV load in leaf tissue and virus acquisition by the thrips vector [55]. In ectoparasites, the number of transmission stages (adult daughters for *T. urticae*) is a direct measure of transmission potential [56] and in line with this we previously demonstrated that higher numbers of *T. urticae* adult daughters correlates positively with transmission [39].

Looking at quantitative relationships between host and parasite traits can enable virulence to be decomposed into exploitation (mean density) and per pathogen pathogenicity (PPP; damage per individual parasite as slope of the relationship between density and harm) [57,58]. It would also be possible for coinfection to impact these different measures of virulence differently. Yet, despite higher viral load and virulence for the France81 isolate, the slope of the relationship between these traits (measured for height, weight and flowers) did not differ between viral isolates and were unaffected by *T. urticae* (see Supplementary Table S7 for these additional analyses). Thus, differences among the TSWV isolates we used were probably due to differences in exploitation, rather than per parasite pathogenicity. Another study identified differences in slopes between viral load and virulence for low versus high viral accumulation phenotypes of cauliflower mosaic virus indicating differences in PPP [47] and variation in exploitation and PPP was found for four different bacteria species infecting *Drosophila melanogaster* [59]. Further, experimental evolution selected for a change in the slope between viral load and virulence in potato virus Y and the relationship changed on different genotypes of host plant [60]. Although not relevant here, these components of virulence may respond differently to coinfection in other systems changing consequences for parasite life-history and evolution.

There was variation in virulence and load between the viral isolates. Overall, plants infected with the France81 isolate had higher systemic viral load, were smaller and had fewer flowers than plants infected with isolate LYE1137vir. Other studies also found variation in the degree of necrosis, stunting [34] and the lesion phenotype [61] induced by different TSWV isolates. Variation in viral traits may arise because of trade-offs among life-history traits [62]. Contrary to France81, TSWV isolate LYE1137vir is able to infect pepper plant varieties carrying the *Tsw* resistance gene. This enlarged host range is conferred by mutations in the TSWV gene that encodes the NSs protein, an inhibitor of general antiviral plant defenses [63]. Consequently, there may be a trade-off between viral load and virulence on one side, and its gain in capacity to infect *Tsw* pepper plants on the other side, due to pleiotropy or linkage between mutations in the NSs gene. Costs of generalism have been identified in other parasites [64,65] and costs associated with overcoming host resistance were found in many plant pathogens, for example in the bacterial blight *Xanthomonas oryzae. pv. oryzae* infecting rice [66] and the wheat fungus *Puccinia striiformis f.sp.tritici* [67].

### Summary

Our results show the importance of measuring the life-history traits of all players, both parasites and the host, when investigating the effects of coinfection on parasite fitness. Our measurements for both TSWV viral load and the number of *T. urticae* are not different in single or coinfection. It is only when we consider virulence that consequences for parasite fitness are revealed. Notably, the most virulent TSWV isolate, France81, enables *T. urticae* to produce equivalent numbers of offspring on much smaller plants. Future work should focus on uncovering how parasite fitness in coinfection changes over the life-span of an entire infection and/or until host death.

## Acknowledgements

We would like to thank Sophie Armitage for the suggestion of decomposing virulence into PPP and exploitation and Yannis Michalakis for the insightful discussions about the results and the project. We would like to thank Sarah Grosjean, and Marie Challe for their help in maintaining the spider mite populations. This is publication n° ISEM 2026-251.

## Data availability statement

Raw data and scripts for the statistical analyses in the manuscript are available at doi:10.5281/zenodo.23090060.

**Supplementary materials** are available in a separate accompanying document found at 10.5281/zenodo.23090060.

