## Supplementary Materials for "Incorporating host traits into analysis of parasite life history reveals facilitation in coinfection experiments" (from another version of this preprint: version 5, 2026-07-01)

Figure S1: Effect of tomato spotted wilt virus (TSWV) infection on mean (± standard error) a) weight, b) number of flowers and chlorophyll content in c) upper and d) lower leaves of plants singly inoculated with isolate France81, LYE1137vir or mock-inoculated in the presence (filled circles) or absence (open circles) of *T. urticae*. All these traits were measured on day 28 after virus inoculation. Small circles refer to results from individual replicate plants.

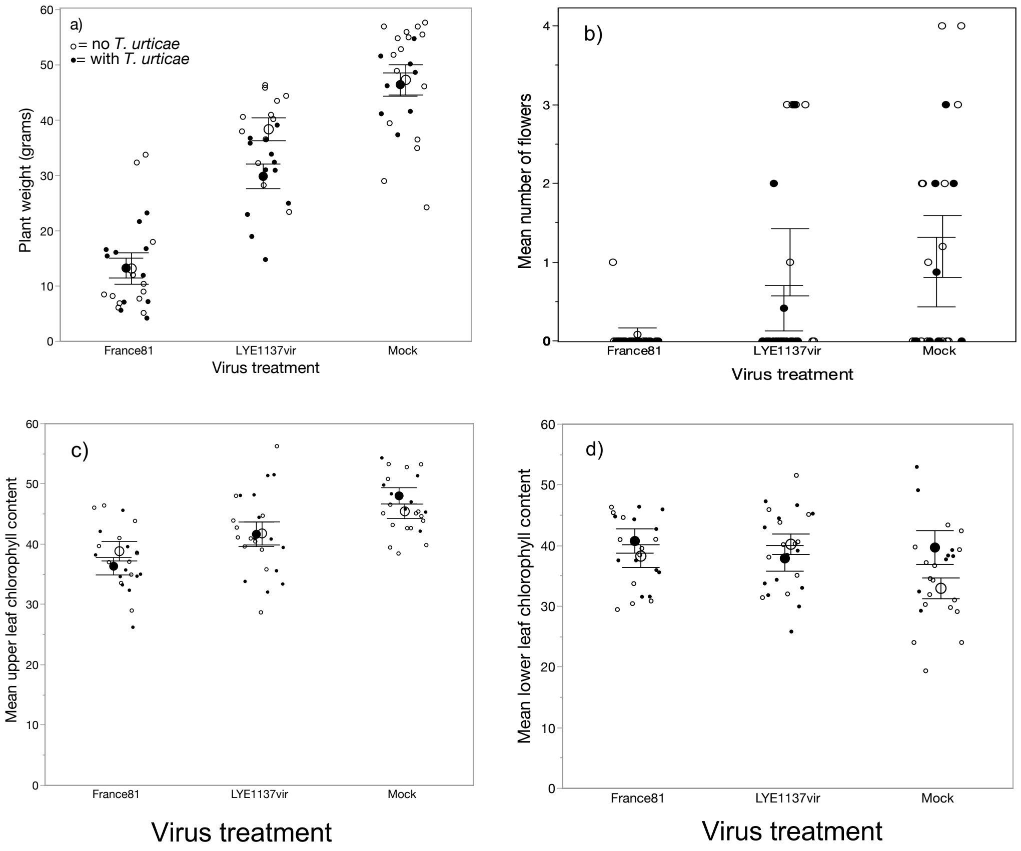

Figure S2: Mean offspring sex ratio (± binomial confidence intervals) on mock-inoculated plants and plants infected with either TSWV isolate France81 or LYE1137vir.

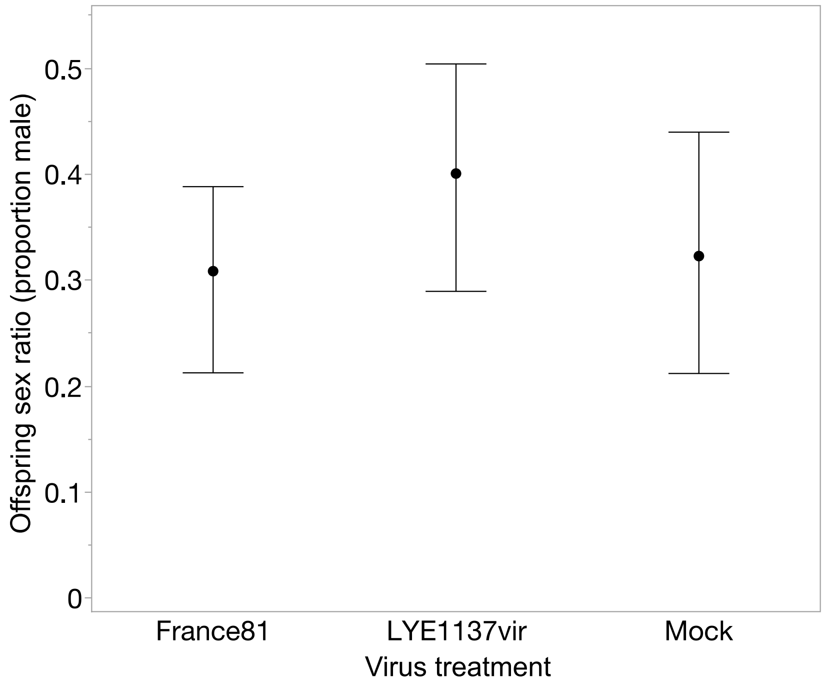

Table S1: Statistical analysis investigating the effects of virus isolate, presence or absence of *T. urticae* mites and the day after infection on the height of singly-infected and coinfected host plants. Note there are two separate general linear mixed models (GLMM)with a Gaussian error structure shown: the first testing how viral isolate versus mock-inoculated plants on days 13, 21 and 27 impacts height, and a second including presence or absence of *T. urticae*, viral isolate, day (21 and 27) and their interactions. The table shows the degrees of freedom (d.f.), the F-ratio, and the associated p-value from the minimal model containing each term. Significant terms are shown in bold. Plant replicate, nested within virus (and mite), was included as a random factor. The proportion of the variance explained by this term for each model (± standard error) is also shown. Significant terms are shown in bold.

| | Plant height in absence of mites on days 13, 21 and 27 | | | Plant height in presence of mites on days 21 and 27 | | |
|---|---|---|---|---|---|---|
| Variable | d.f. | F-ratio | p-value | d.f. | F-ratio | p-value |
| Virus | **2, 85** | **89.11** | **< 0.001** | **2, 109** | **108.84** | **< 0.001** |
| Mites | | | | **1, 111** | **12.10** | **0.0007** |
| Day | **2, 102** | **105.49** | **< 0.001** | **1, 122** | **79.56** | **< 0.001** |
| Virus*Mite | | | | 2, 107 | 2.80 | 0.0653 |
| Virus*Day | **4, 106** | **6.47** | **< 0.001** | **2, 126** | **5.73** | **0.0042** |
| Mite*Day | | | | 1, 28 | 0.38 | 0.5400 |
| Virus*Mite*Day | | | | 2, 122 | 0.32 | 0.7267 |
| Plant replicate | **17.44 ± 3.40** | | | **20.01 ± 3.04** | | |

Table S2: Statistical analysis of general linear models (GLM) investigating the effects of virus isolate and mite presence or absence on tomato plant weight, number of flowers, and chlorophyll content of upper and lower leaves. All measures were taken on day 28 after virus inoculation. The error distribution attributed in each model is stated in brackets after the variable name. The results shown (d.f. = degrees of freedom, $\chi^2$= the chi-square values and p-values) are from the minimal model containing each term. Significant terms are shown in bold.

| | **Plant weight (Gaussian)** | | | **Number of Flowers (Poisson)** | | | **Chlorophyll (upper leaves) (Gaussian)** | | | **Chlorophyll (lower leaves) (Gaussian)** | | |
|---|---|---|---|---|---|---|---|---|---|---|---|---|
| Variable | d.f. | $\chi^2$ | p-value | d.f. | $\chi^2$ | p-value | d.f. | $\chi^2$ | p-value | d.f. | Chisq | p-value |
| Virus | **2** | **93.86** | **<0.001** | **2** | **17.20** | **0.0002** | **2** | **25.31** | **<0.001** | 2 | 5.55 | 0.0623 |
| Mites | 1 | 2.47 | 0.1160 | 1 | 2.10 | 0.1477 | 1 | 0.004 | 0.9493 | 1 | 1.65 | 0.1988 |
| Virus* Mite | 2 | 3.88 | 0.1440 | 2 | 1.00 | 0.6078 | 2 | 2.42 | 0.2975 | 2 | 5.26 | 0.0722 |

Table S3: Statistical analysis of GLMs with a Gaussian error distribution investigating the effect of virus isolate, presence or absence of *T. urticae* mites, and the day post inoculation on titres of France81 and LYE1137vir TSWV isolates. The first model shows viral titre in plants singly infected by the virus on days 7, 13, 21 and 27 and the second model shows viral titre in plants coinfected with *T. urticae* on days 21 and 27 only. The table shows the degrees of freedom (d.f.), the $\chi^2$, and the associated p-value from the minimal model containing each term. Significant terms are shown in bold.

| | **Virus single infection Days 7, 13, 20 and 27** | | | **Virus and mite coinfection Days 21 and 27** | | |
|---|---|---|---|---|---|---|
| Variable | d.f. | $\chi^2$ | p-value | d.f. | $\chi^2$ | p-value |
| Virus | **1** | **17.60** | **<0.001** | **1** | **11.82** | **<0.001** |
| Day | **3** | **112.37** | **<0.001** | **1** | **11.94** | **<0.001** |
| Mite | | | | 1 | 0.26 | 0.6105 |
| Day*Virus | 3 | 1.59 | 0.6627 | 1 | 2.27 | 0.1321 |
| Day*Mite | | | | 1 | 0.26 | 0.6111 |
| Virus*Mite | | | | 1 | 0.21 | 0.6423 |
| Day*Virus*Mite | | | | 1 | 0.18 | 0.8924 |

Table S4: Statistical analysis of GLMMs and a GLM showing the effect of virus isolate, leaf sampled and their interaction on the total number of adult females and virus isolate on offspring sex ratio and number of adult daughters per cm of plant height. The type of model and error distribution are shown in brackets after the response variable. The table shows the degrees of freedom (d.f.), the F-ratio, and the associated p-value from the minimal model containing each term. Models for the total number of adult females and sex ratio included plant replicate, and all models included fungus presence/absence, as random factors and the proportions of variance ± standard error for each are shown. Significant terms are shown in bold.

| | **Total number of adult daughters (GLMM, negative binomial)** | | | **Total number of daughters per cm (GLMM, normal)** | | | **Sex ratio (GLM, binomial)** | | |
|---|---|---|---|---|---|---|---|---|---|
| Variable | d.f. | F-ratio | p-value | d.f. | F-ratio | p-value | d.f. | F-ratio | p-value |
| Virus | 2, 27 | 0.76 | 0.4769 | **2, 19** | **18.72** | **<0.0.001** | 2, 15 | 1.08 | 0.3646 |
| Leaf | **1, 29** | **4.76** | **0.0375** | | | | | | |
| Virus *leaf | 2, 27 | 0.06 | 0.9396 | | | | | | |
| Fungus | 0.09 ± 0.15 | | | 1.58 ± 2.56 | | | 0.01 ± 0.02 | | |
| Plant ID | 0.06 ± 0.09 | | | | | | 0.22 ± 0.9 | | |

Table S5: Eigenvalues and percentages of explained variance for the PCA of seven traits measured on host plant, TSWV and *T. urticae*.

| | PC1 | PC2 | PC3 | PC4 | PC5 | PC6 | PC7 |
|---|---|---|---|---|---|---|---|
| Eigenvalue | 2.78 | 1.38 | 1.03 | 0.85 | 0.65 | 0.24 | 0.07 |
| Percentage of explained variance | 39.71 | 19.82 | 14.72 | 12.10 | 9.27 | 3.36 | 1.03 |
| Cumulative percentage of explained variance | 39.71 | 59.53 | 74.25 | 86.33 | 95.60 | 98.96 | 100 |

Table S6: Loading values of the PCA for the seven host, TSWV and *T. urticae* traits.

| | | PC1 | PC2 | PC3 | PC4 | PC5 | PC6 | PC7 |
|---|---|---|---|---|---|---|---|---|
| Host | Height | 0.9271 | 0.1094 | -0.2516 | 0.1213 | -0.0067 | -0.0979 | 0.2023 |
| | Weight | 0.9401 | 0.0482 | -0.0512 | 0.1205 | -0.0435 | -0.2610 | -0.1637 |
| | Flowers | 0.4938 | 0.3375 | 0.3318 | -0.5750 | 0.4400 | 0.0890 | 0.0050 |
| | Upper leaf chlorophyll | 0.6141 | -0.4477 | 0.2985 | 0.4751 | 0.1864 | 0.2693 | -0.0183 |
| | Lower leaf chlorophyll | -0.0046 | 0.5008 | 0.7802 | 0.2093 | -0.3000 | -0.0760 | 0.0316 |
| Virus | Log relative load | -0.6168 | 0.3141 | -0.0191 | 0.4301 | 0.5629 | -0.1350 | 0.0231 |
| *T. urticae* | Female offspring | 0.1870 | 0.8422 | -0.3954 | 0.1755 | -0.1103 | 0.2307 | -0.0560 |

Table S7: Statistical analysis of GLMMs with a Gaussian error structure, and a GLM with a Poisson error structure, investigating the effects of virus isolate, mite coinfection status and viral titre on the height and weight of host plants on days 21 and 28 post inoculation and the number of flowers on day 28. The results shown (d.f. = degrees of freedom, F ratios or $\chi^2$ and p-value) are from the minimal model containing each term. Significant terms are shown in bold. The proportion of plant height and weight variance explained by the day variable (± standard error) is also shown.

| | **Plant height (Gaussian)** | | | **Plant weight (Gaussian)** | | | **Number of flowers (Poisson)** | | |
|---|---|---|---|---|---|---|---|---|---|
| Variable | d.f. | F-ratio | p-value | d.f. | F-ratio | p-value | d.f. | $\chi^2$ | p-value |
| Virus | **1, 91** | **79.71** | **<0.0001** | **1, 91** | **90.76** | **< 0.0001** | **1** | **10.54** | **0.0012** |
| Mite | **1, 91** | **10.27** | **0.0019** | 1, 91 | 1.30 | 0.2563 | 1 | 2.45 | 0.1153 |
| Virus titre | 1, 79 | 0.24 | 0.6276 | 1, 79 | 0.73 | 0.3948 | 1 | 0.37 | 0.5445 |
| Virus*Mite | 1, 90 | 3.56 | 0.0617 | **1, 91** | **5.12** | **0.0262** | 1 | 0.32 | 0.5738 |
| Virus*Virus titre | 1, 77 | 0.36 | 0.5472 | 1, 78 | 3.70 | 0.0581 | 1 | 0.02 | 0.8984 |
| Virus titre*Mite | 1, 76 | 0.02 | 0.8768 | 1, 76 | 0.09 | 0.7680 | 1 | 3.59 | 0.0582 |
| Viral titre*Virus*Mite | 1, 75 | 0.27 | 0.6078 | 1, 75 | 0.05 | 0.8246 | 1 | 0.000 | 0.9998 |
| **Day** | 20.32 ± 29.57 | | | 23.01 ± 35.00 | | | | | |

*Description of statistical analyses in Table S7*

To get a more thorough understanding of relationships among traits, we used a GLMM with a normal error distribution to see if plant height and weight was affected by viral load for virus-infected plants on days 20 and 27. We used a GLM with a Poisson distribution to see if viral load affected the number of flowers. In these models, log relative viral concentration was included as a covariate, viral isolate and spider mite absence and presence as fixed factors and their interactions. Day of sampling was included as a random factor in the models for weight and height.

*Summary of results in Table S7*

We modelled the effect of viral titre on plant traits (height, weight and flowers) in the presence or absence of *T. urticae* for the different TSWV isolates (excluding replicates from the mock-inoculation treatment) to investigate different components of virulence, notably proliferation and per parasite pathogenicity. These models were in accordance with the results found in earlier models that viral isolate impacted plant height ($F_{1, 91} = 79.71$, $p < 0.0001$), weight ($F_{1, 77} = 58.65$, $p < 0.0001$), and flowers ($\chi^2 = 10.54$, $p = 0.0012$) with isolate France81 reducing these traits more than isolate LYE137vir, and that the presence of *T. urticae* reduced plant height ($F_{1, 91} = 10.27$, $p = 0.0019$), but not flowers ($\chi^2 = 2.48$, $p = 0.1153$) (Tables S1 and S7 for full models). Although the main effect for *T. urticae* was not significant ($F_{1, 77} = 0.92$, $p = 0.3411$) there was a significant interaction between viral isolate and mite ($F_{1, 77} = 7.73$, $p = 0.0068$) showing that plants coinfected with isolate LYE1137vir and *T. urticae* were lighter than plants singly infected with the LYE1137vir isolate. We were, however, mostly interested in whether the main effect for viral titre or its interactions with other explanatory variables explained a significant amount of the variance in these models (Table S7). There was no significant main effect of viral titre for plant height ($F_{1, 79} = 0.24$, $p = 0.6267$), weight ($F_{1, 78} = 0.02$, $p = 0.9001$) or flowers ($\chi^2 = 0.37$, $p = 0.5445$). Further, there was no significant interaction between viral titre and other traits for height, weight and flowers.
