## Supplementary material for "Incorporating host traits into analysis of parasite life history reveals facilitation in coinfection experiments": Script for statistical analyses (from another version of this preprint: version 5, 2026-07-01)

**Script for analyses in JMP**

***1. Plant traits:*** Model results are shown in Tables S1 and S2. Full models are shown followed by simplified versions removing significant terms in a stepwise fashion for terms with p-values >0.05 (if applicable).

1.1. General linear mixed model (GLMM) with a Gaussian error structure for plant height for mite free plants on days 13, 21 and 27.

```
Fit Model(
	Y( :Height ),
	Effects( :Virus, :Day, :Virus * :Day ),
	Random Effects( :Data sheet plant number[:Virus] ),
	NoBounds( 1 ),
	Personality( "Generalized Linear Mixed Model" ),
	Generalized Distribution( "Normal" ),
	Run( Fit )
);
```

1.2. GLMMs with a Gaussian error structure investigating the effect of mite, virus, day and their interactions on plant height on days 21 and 27.

```
Fit Model(
	Y( :Height ),
	Effects(
		:Virus, :Mite, :Virus * :Mite, :Day, :Virus * :Day, :Mite * :Day,
		:Virus * :Mite * :Day
	),
	Random Effects( :Data sheet plant number[:Virus, :Mite] ),
	NoBounds( 1 ),
	Personality( "Generalized Linear Mixed Model" ),
	Generalized Distribution( "Normal" ),
	Run( Fit )
);

Fit Model(
	Y( :Height ),
	Effects( :Virus, :Mite, :Virus * :Mite, :Day, :Virus * :Day, :Mite * :Day ),
	Random Effects( :Data sheet plant number[:Virus, :Mite] ),
	NoBounds( 1 ),
	Personality( "Generalized Linear Mixed Model" ),
	Generalized Distribution( "Normal" ),
	Run( Fit )
);

Fit Model(
	Y( :Height ),
	Effects( :Virus, :Mite, :Virus * :Mite, :Day, :Virus * :Day ),
	Random Effects( :Data sheet plant number[:Virus, :Mite] ),
	NoBounds( 1 ),
	Personality( "Generalized Linear Mixed Model" ),
	Generalized Distribution( "Normal" ),
```

```
        Run( Fit )
);


Fit Model(
        Y( :Height ),
        Effects( :Virus, :Mite, :Day, :Virus * :Day ),
        Random Effects( :Data sheet plant number[:Virus, :Mite] ),
        NoBounds( 1 ),
        Personality( "Generalized Linear Mixed Model" ),
        Generalized Distribution( "Normal" ),
        Run( Fit )
);
```

1.3. GLMs with a Gaussian error structure investigating the effect of mite, virus and their interactions on plant weight on day 28.

```
Fit Model(
        Y( :Weight ),
        Effects( :Virus, :Mite, :Virus * :Mite ),
        Personality( "Generalized Linear Model" ),
        GLM Distribution( "Normal" ),
        Link Function( "Identity" ),
        Overdispersion Tests and Intervals( 0 ),
        "Firth Bias-Adjusted Estimates"n( 0 ),
        Run
);

Fit Model(
        Y( :Weight ),
        Effects( :Virus, :Mite ),
        Personality( "Generalized Linear Model" ),
        GLM Distribution( "Normal" ),
        Link Function( "Identity" ),
        Overdispersion Tests and Intervals( 0 ),
        "Firth Bias-Adjusted Estimates"n( 0 ),
        Run
);

Fit Model(
        Y( :Weight ),
        Effects( :Virus ),
        Personality( "Generalized Linear Model" ),
        GLM Distribution( "Normal" ),
        Link Function( "Identity" ),
        Overdispersion Tests and Intervals( 0 ),
        "Firth Bias-Adjusted Estimates"n( 0 ),
        Run
);
```

1.4. GLMs with a Poisson error structure investigating the effect of mite, virus and their interaction on plant weight on day 28.

```
Fit Model(
	Y( :Flowers ),
	Effects( :Virus, :Mite, :Virus * :Mite ),
	Personality( "Generalized Linear Model" ),
	GLM Distribution( "Poisson" ),
	Link Function( "Log" ),
	Overdispersion Tests and Intervals( 1 ),
	"Firth Bias-Adjusted Estimates"n( 0 ),
	Run
);

Fit Model(
	Y( :Flowers ),
	Effects( :Virus, :Mite ),
	Personality( "Generalized Linear Model" ),
	GLM Distribution( "Poisson" ),
	Link Function( "Log" ),
	Overdispersion Tests and Intervals( 1 ),
	"Firth Bias-Adjusted Estimates"n( 0 ),
	Run
);

Fit Model(
	Y( :Flowers ),
	Effects( :Virus ),
	Personality( "Generalized Linear Model" ),
	GLM Distribution( "Poisson" ),
	Link Function( "Log" ),
	Overdispersion Tests and Intervals( 1 ),
	"Firth Bias-Adjusted Estimates"n( 0 ),
	Run
);
```

1.5. GLMs with a Gaussian error structure investigating the effect of mite, virus and their interaction on upper leaf chlorophyll content on day 28.

```
Fit Model(
	Y( :upper mean ),
	Effects( :Virus, :Mite, :Virus * :Mite ),
	Personality( "Generalized Linear Model" ),
	GLM Distribution( "Normal" ),
	Link Function( "Identity" ),
	Overdispersion Tests and Intervals( 0 ),
	"Firth Bias-Adjusted Estimates"n( 0 ),
	Run
);

Fit Model(
```

```
        Y( :upper mean ),
        Effects( :Virus, :Mite ),
        Personality( "Generalized Linear Model" ),
        GLM Distribution( "Normal" ),
        Link Function( "Identity" ),
        Overdispersion Tests and Intervals( 0 ),
        "Firth Bias-Adjusted Estimates"n( 0 ),
        Run
);

Fit Model(
        Y( :upper mean ),
        Effects( :Virus ),
        Personality( "Generalized Linear Model" ),
        GLM Distribution( "Normal" ),
        Link Function( "Identity" ),
        Overdispersion Tests and Intervals( 0 ),
        "Firth Bias-Adjusted Estimates"n( 0 ),
        Run
);
```

1.6. GLMs with a Gaussian error structure investigating the effect of mite, virus and their interaction on lower leaf chlorophyll content on day 28.

```
Fit Model(

        Y( :Lower mean ),
        Effects( :Virus, :Mite, :Virus * :Mite ),
        Personality( "Generalized Linear Model" ),
        GLM Distribution( "Normal" ),
        Link Function( "Identity" ),
        Overdispersion Tests and Intervals( 0 ),
        "Firth Bias-Adjusted Estimates"n( 0 ),
        Run
);

Fit Model(
        Y( :Lower mean ),
        Effects( :Virus, :Mite ),
        Personality( "Generalized Linear Model" ),
        GLM Distribution( "Normal" ),
        Link Function( "Identity" ),
        Overdispersion Tests and Intervals( 0 ),
        "Firth Bias-Adjusted Estimates"n( 0 ),
        Run
);

Fit Model(
        Y( :Lower mean ),
        Effects( :Virus ),
        Personality( "Generalized Linear Model" ),
        GLM Distribution( "Normal" ),
```

```
        Link Function( "Identity" ),
        Overdispersion Tests and Intervals( 0 ),
        "Firth Bias-Adjusted Estimates"n( 0 ),
        Run
);
```

***2. TSWV traits***. The outputs for these models are shown in Table S3. The full models are shown for each trait, followed by models that were simplified by removing significant terms in a stepwise fashion for terms with p-values >0.05.

2.1. GLMs with a Gaussian error structure investigating the effect of virus isolate, day and their interaction on log relative TSWV concentration for mite free plants on days 13, 21 and 27.

```
Fit Model(
        Y( :log rel concentration ),
        Effects( :Virus, :Day, :Virus * :Day ),
        Personality( "Generalized Linear Model" ),
        GLM Distribution( "Normal" ),
        Link Function( "Identity" ),
        Overdispersion Tests and Intervals( 0 ),
        "Firth Bias-Adjusted Estimates"n( 0 ),
        Run
);

Fit Model(
        Y( :log rel concentration ),
        Effects( :Virus, :Day ),
        Personality( "Generalized Linear Model" ),
        GLM Distribution( "Normal" ),
        Link Function( "Identity" ),
        Overdispersion Tests and Intervals( 0 ),
        "Firth Bias-Adjusted Estimates"n( 0 ),
        Run
);
```

2.2 GLMs with a Gaussian error structure investigating the effect of virus isolate, mite presence and day and their interactions on log relative TSWV concentration for mite free plants on days 21 and 27.

```
Fit Model(
        Y( :log rel concentration ),
        Effects(
                :Virus, :Mite, :Virus * :Mite, :Day, :Virus * :Day, :Mite * :Day,
                :Virus * :Mite * :Day
        ),
        Personality( "Generalized Linear Model" ),
        GLM Distribution( "Normal" ),
        Link Function( "Identity" ),
```

```
		Overdispersion Tests and Intervals( 0 ),
		"Firth Bias-Adjusted Estimates"n( 0 ),
		Run
);

Fit Model(
		Y( :log rel concentration ),
		Effects( :Virus, :Mite, :Virus * :Mite, :Day, :Virus * :Day, :Mite * :Day ),
		Personality( "Generalized Linear Model" ),
		GLM Distribution( "Normal" ),
		Link Function( "Identity" ),
		Overdispersion Tests and Intervals( 0 ),
		"Firth Bias-Adjusted Estimates"n( 0 ),
		Run
);

Fit Model(
		Y( :log rel concentration ),
		Effects( :Virus, :Mite, :Day, :Virus * :Day, :Mite * :Day ),
		Personality( "Generalized Linear Model" ),
		GLM Distribution( "Normal" ),
		Link Function( "Identity" ),
		Overdispersion Tests and Intervals( 0 ),
		"Firth Bias-Adjusted Estimates"n( 0 ),
		Run
);

Fit Model(
		Y( :log rel concentration ),
		Effects( :Virus, :Mite, :Day, :Virus * :Day ),
		Personality( "Generalized Linear Model" ),
		GLM Distribution( "Normal" ),
		Link Function( "Identity" ),
		Overdispersion Tests and Intervals( 0 ),
		"Firth Bias-Adjusted Estimates"n( 0 ),
		Run
);

Fit Model(
		Y( :log rel concentration ),
		Effects( :Virus, :Mite, :Day ),
		Personality( "Generalized Linear Model" ),
		GLM Distribution( "Normal" ),
		Link Function( "Identity" ),
		Overdispersion Tests and Intervals( 0 ),
		"Firth Bias-Adjusted Estimates"n( 0 ),
		Run
);

Fit Model(
		Y( :log rel concentration ),
```

```
        Effects( :Virus, :Day ),
        Personality( "Generalized Linear Model" ),
        GLM Distribution( "Normal" ),
        Link Function( "Identity" ),
        Overdispersion Tests and Intervals( 0 ),
        "Firth Bias-Adjusted Estimates"n( 0 ),
        Run
);
```

***3. T. urticae traits;*** The outputs for these models are shown in Table S4. The full models are shown for each trait, followed by models that were simplified by removing significant terms in a stepwise fashion for terms with p-values >0.05 (if applicable).

3.1. GLMM with a negative binomial error structure investigating the effect of virus isolate, leaf level and the interaction between them on total number of *T. urticae* adult daughters. The data for number of daughters on leaf 1 and leaf 2 were stacked for this analysis and a new column 'Leaf level' created. The presence of fungus detected on plants was included in the model as a random effect. Plant identity was also included in the model as a random term to control for overdispersion.

```
Fit Model(
        Y( :Data ),
        Effects( :Virus, :Leaf level, :Virus * :Leaf level ),
        Random Effects( :Fungus, :ID ),
        NoBounds( 1 ),
        Personality( "Generalized Linear Mixed Model" ),
        Generalized Distribution( "Negative Binomial" ),
        Run( Fit )
);
Fit Model(
        Y( :Data ),
        Effects( :Virus, :Leaf level ),
        Random Effects( :Fungus, :ID ),
        NoBounds( 1 ),
        Personality( "Generalized Linear Mixed Model" ),
        Generalized Distribution( "Negative Binomial" ),
        Run( Fit )
);
Fit Model(
        Y( :Data ),
        Effects( :Leaf level ),
        Random Effects( :Fungus, :ID ),
        NoBounds( 1 ),
        Personality( "Generalized Linear Mixed Model" ),
        Generalized Distribution( "Negative Binomial" ),
        Run( Fit )
);
```

3.2. GLMM with a binomial error structure investigating the effect of viral isolate on *T. urticae* offspring sex ratio. The presence of fungus detected on plants was included in the model as a random effect. Plant identity was also included in the model as a random term to control for overdispersion.

```
Fit Model(
	Y( :Total males, :Total Adults ),
	Effects( :Virus ),
	Random Effects( :Fungus, :ID ),
	NoBounds( 1 ),
	Personality( "Generalized Linear Mixed Model" ),
	Generalized Distribution( "Binomial" ),
	Link Function( "Logit" ),
	Run( Fit )
);
```

3.3: GLMM investigating the effect of viral isolate on *T. urticae* females per cm of plant height. The presence of fungus detected on plants was included in the model as a random effect.

```
Fit Model(
	Y( :Females per cm ),
	Effects( :Virus ),
	Random Effects( :Fungus ),
	Keep dialog open( 1 ),
	Personality( "Generalized Linear Mixed Model" ),
	Generalized Distribution( "Normal" )
);
```

3.4: Principal Components Analysis on height, virus concentration, weight, flowers upper leaf mean chlorophyll content, lower mean chlorophyll content and total adult daughters. Model outputs are shown in Tables S5 and S6.

```
Principal Components(
	Y(
		:log rel concentration, :Height, :Weight, :Flowers, :upper mean, :Lower mean,
		:Total females
	),
	Estimation Method( "REML" ),
	Standardize( "Standardized" ),
	Correlations( 1 ),
	Biplot( 2 )
);
```

***4. Models investigating how virus concentration impacts host traits.*** The outputs for these models are shown in Table S7. The full models are shown for each trait, followed by models that were simplified by removing significant terms in a stepwise fashion for terms with p-values >0.05.

4.1. GLMM with a Gaussian error structure investigating the effect of virus, virus concentration, mite and their interactions on plant height tolerance on days 21 and 27. Day was included in models as a random effect.

```
Fit Model(
	Y( :Height ),
	Effects(
		:Virus, :Mite, :Virus * :Mite, :log rel concentration,
		:Virus * :log rel concentration, :Mite * :log rel concentration,
		:Virus * :Mite * :log rel concentration
	),
	Random Effects( :Day ),
	NoBounds( 1 ),
	Personality( "Generalized Linear Mixed Model" ),
	Generalized Distribution( "Normal" ),
	Run( Fit )
);

Fit Model(
	Y( :Height ),
	Effects(
		:Virus, :Mite, :Virus * :Mite, :log rel concentration,
		:Virus * :log rel concentration, :Mite * :log rel concentration
	),
	Random Effects( :Day ),
	NoBounds( 1 ),
	Personality( "Generalized Linear Mixed Model" ),
	Generalized Distribution( "Normal" ),
	Run( Fit )
);

Fit Model(
	Y( :Height ),
	Effects(
		:Virus, :Mite, :Virus * :Mite, :log rel concentration,
		:Virus * :log rel concentration
	),
	Random Effects( :Day ),
	NoBounds( 1 ),
	Personality( "Generalized Linear Mixed Model" ),
	Generalized Distribution( "Normal" ),
	Run( Fit )
);

Fit Model(
	Y( :Height ),
	Effects( :Virus, :Mite, :Virus * :Mite, :log rel concentration ),
	Random Effects( :Day ),
	NoBounds( 1 ),
	Personality( "Generalized Linear Mixed Model" ),
	Generalized Distribution( "Normal" ),
```

```
	Run( Fit )
);

Fit Model(
	Y( :Height ),
	Effects( :Virus, :Mite, :Virus * :Mite ),
	Random Effects( :Day ),
	NoBounds( 1 ),
	Personality( "Generalized Linear Mixed Model" ),
	Generalized Distribution( "Normal" ),
	Run( Fit )
);

Fit Model(
	Y( :Height ),
	Effects( :Virus, :Mite ),
	Random Effects( :Day ),
	NoBounds( 1 ),
	Personality( "Generalized Linear Mixed Model" ),
	Generalized Distribution( "Normal" ),
	Run( Fit )
);
```

4.2. GLMM with a Gaussian error structure investigating the effect of virus, virus concentration, mite and their interactions on plant weight on days 21 and 27. Day was included in models as a random effect.

```
Fit Model(
	Y( :Weight ),
	Effects(
		:Virus, :Mite, :Virus * :Mite, :log rel concentration,
		:Virus * :log rel concentration, :Mite * :log rel concentration,
		:Virus * :Mite * :log rel concentration
	),
	Random Effects( :Day ),
	NoBounds( 1 ),
	Personality( "Generalized Linear Mixed Model" ),
	Generalized Distribution( "Normal" ),
	Run( Fit )
);

Fit Model(
	Y( :Weight ),
	Effects(
		:Virus, :Mite, :Virus * :Mite, :log rel concentration,
		:Virus * :log rel concentration, :Mite * :log rel concentration
	),
	Random Effects( :Day ),
	NoBounds( 1 ),
	Personality( "Generalized Linear Mixed Model" ),
```

```
		Generalized Distribution( "Normal" ),
		Run( Fit )
);

Fit Model(
		Y( :Weight ),
		Effects(
				:Virus, :Mite, :Virus * :Mite, :log rel concentration,
				:log rel concentration * :Virus
		),
		Random Effects( :Day ),
		NoBounds( 1 ),
		Personality( "Generalized Linear Mixed Model" ),
		Generalized Distribution( "Normal" ),
		Run( Fit )
);

Fit Model(
		Y( :Weight ),
		Effects( :Virus, :Mite, :Virus * :Mite, :log rel concentration ),
		Random Effects( :Day ),
		NoBounds( 1 ),
		Personality( "Generalized Linear Mixed Model" ),
		Generalized Distribution( "Normal" ),
		Run( Fit )
);

Fit Model(
		Y( :Weight ),
		Effects( :Virus, :Mite, :Virus * :Mite ),
		Random Effects( :Day ),
		NoBounds( 1 ),
		Personality( "Generalized Linear Mixed Model" ),
		Generalized Distribution( "Normal" ),
		Run( Fit )
);
```

4.3. GLM with a Poisson error structure investigating the effect of virus, virus concentration, mite and their interactions on the number of flowers on day 27.

```
Fit Model(
		Y( :Flowers ),
		Effects(
				:Virus, :Mite, :Virus * :Mite, :log rel concentration,
				:Virus * :log rel concentration, :Mite * :log rel concentration,
				:Virus * :Mite * :log rel concentration
		),
		Personality( "Generalized Linear Model" ),
		GLM Distribution( "Poisson" ),
		Link Function( "Log" ),
		Overdispersion Tests and Intervals( 1 ),
```

```
		"Firth Bias-Adjusted Estimates"n( 0 ),
		Run
);

Fit Model(
		Y( :Flowers ),
		Effects(
			:Virus, :Mite, :Virus * :Mite, :log rel concentration,
			:Virus * :log rel concentration, :Mite * :log rel concentration
		),
		Personality( "Generalized Linear Model" ),
		GLM Distribution( "Poisson" ),
		Link Function( "Log" ),
		Overdispersion Tests and Intervals( 1 ),
		"Firth Bias-Adjusted Estimates"n( 0 ),
		Run
);

Fit Model(
		Y( :Flowers ),
		Effects(
			:Virus, :Mite, :Virus * :Mite, :log rel concentration,
			:Mite * :log rel concentration
		),
		Personality( "Generalized Linear Model" ),
		GLM Distribution( "Poisson" ),
		Link Function( "Log" ),
		Overdispersion Tests and Intervals( 1 ),
		"Firth Bias-Adjusted Estimates"n( 0 ),
		Run
);

Fit Model(
		Y( :Flowers ),
		Effects( :Virus, :Mite, :log rel concentration, :Mite * :log rel concentration ),
		Personality( "Generalized Linear Model" ),
		GLM Distribution( "Poisson" ),
		Link Function( "Log" ),
		Overdispersion Tests and Intervals( 1 ),
		"Firth Bias-Adjusted Estimates"n( 0 ),
		Run
);

Fit Model(
		Y( :Flowers ),
		Effects( :Virus, :Mite, :log rel concentration ),
		Personality( "Generalized Linear Model" ),
		GLM Distribution( "Poisson" ),
		Link Function( "Log" ),
		Overdispersion Tests and Intervals( 1 ),
		"Firth Bias-Adjusted Estimates"n( 0 ),
```

```
	Run
);

Fit Model(
	Y( :Flowers ),
	Effects( :Virus, :Mite ),
	Personality( "Generalized Linear Model" ),
	GLM Distribution( "Poisson" ),
	Link Function( "Log" ),
	Overdispersion Tests and Intervals( 1 ),
	"Firth Bias-Adjusted Estimates"n( 0 ),
	Run
);

Fit Model(
	Y( :Flowers ),
	Effects( :Virus, ),
	Personality( "Generalized Linear Model" ),
	GLM Distribution( "Poisson" ),
	Link Function( "Log" ),
	Overdispersion Tests and Intervals( 1 ),
	"Firth Bias-Adjusted Estimates"n( 0 ),
	Run
);
```
